# Single-molecule imaging reveals translation of mRNAs localized to stress granules

**DOI:** 10.1101/2020.03.31.018093

**Authors:** Daniel Mateju, Bastian Eichenberger, Jan Eglinger, Gregory Roth, Jeffrey A. Chao

## Abstract

Cellular stress leads to reprogramming of mRNA translation and formation of stress granules (SGs), membraneless organelles consisting of mRNA and RNA-binding proteins. Although the function of SGs remains largely unknown, it is widely assumed they contain exclusively nontranslating mRNA. Here we re-examine this hypothesis using single-molecule imaging of mRNA translation in living cells. While our data confirms that non-translating mRNAs are preferentially recruited to SGs, we find unequivocal evidence for translation of mRNA localized to SGs. Our data indicate that SG-associated translation is not rare and that the entire translation cycle (initiation, elongation and termination) can occur for these transcripts. Furthermore, translating mRNAs can be observed transitioning between the cytosol and SGs without changing their translational status. Together, these results argue against a direct role for SGs in inhibition of mRNA translation.

## INTRODUCTION

Cells are often exposed to stress conditions such as oxidative stress, temperature changes, viral infection, or presence of toxins. To protect against these unfavourable conditions and minimize the damage during stress, cells activate an evolutionarily conserved pathway, termed the integrated stress response (Pakos-Zebrucka et al., 2016). This pathway triggers the phosphorylation of the translation initiation factor eIF2α, leading to inhibition of translation of most mRNAs and preferential translation of selected mRNAs such as the transcription factor ATF4. This translational reprogramming allows cells to conserve energy and shift their resources towards the restoration of cellular homeostasis (Pakos-Zebrucka et al., 2016). Along with the reprogramming of translation, the integrated stress response also leads to the assembly of stress granules (SGs), phase-separated membraneless organelles, in the cytoplasm. A fraction of cytoplasmic mRNAs localize to SGs upon their assembly, and translation inhibition and RNA length have been shown to correlate with SG association (Khong et al., 2017). In addition to mRNAs, SGs are enriched in RNA-binding proteins and translation initiation factors (Nunes et al., 2019). There has been significant effort towards understanding SG composition, material properties, and role in diseases, however, their biological function remains largely unknown (Alberti and Dormann, 2019; Guzikowski et al., 2019; Wolozin and Ivanov, 2019).

Recent studies using single-molecule imaging techniques in living cells have begun to shed light on the kinetics of mRNA recruitment to SGs (Moon et al., 2019; Wilbertz et al., 2019). Despite this progress, we know little about the functional consequence of these interactions, as it remains unclear whether sequestration of mRNAs to SGs has any impact on gene expression (Guzikowski et al., 2019; Wilbertz et al., 2019). As SG formation coincides with global silencing of translation, it is often speculated that SGs are directly contributing to this repression during stress (Adjibade et al., 2015; Anderson and Kedersha, 2009; El-Naggar and Sorensen, 2018). This stems from the assumption that SGs contain exclusively nontranslating mRNA and help to keep them in an inhibited state. There is, however, no direct evidence supporting this model because single-molecule methods for measuring the translation and location of transcripts within individual cells have only recently been described (Halstead et al., 2015; Morisaki et al., 2016; Pichon et al., 2016; Wang et al., 2016; Wu et al., 2016; Yan et al., 2016). Furthermore, previous studies have characterized transcripts that are translationally inhibited during stress, making it difficult to distinguish the specific contribution of SGs to this process (Moon et al., 2019; Wilbertz et al., 2019).

While most transcripts are repressed during stress, there are transcripts (e.g. ATF4) that are necessary for mounting the stress response and are preferentially translated only when elF2α is phosphorylated (Harding et al., 2000). In order to characterize the relationship between translational status and SG recruitment of mRNAs during stress, we generated ATF4-SunTag reporter transcripts for single-molecule imaging of translation in living cells. Using this approach, we find direct evidence for active translation of mRNAs localized to SGs. Transcripts localized to SGs can initiate translation, elongate and terminate, indicating that the SG environment does not repress translation. Translating mRNAs can also be observed to dynamically exchange between the cytosol and SGs. Additionally, we show that SG-associated translation is not limited to transcripts whose translation is enhanced during stress, but our measurements are sensitive enough to detect these rare events also for transcripts that are inhibited during stress. As membrane-less organelles are now proposed to play a key role in cellular compartmentalization, these experiments highlight the importance of direct observation of biochemical reactions inside and outside of biomolecular condensates in order to understand their biological function.

## RESULTS

### Single-molecule imaging reveals SG-associated translation

In order to characterize the relationship between mRNA localization and translation during stress, we combined live-cell imaging of SGs with SunTag-based single-molecule imaging of translation (Pichon et al., 2016; Wang et al., 2016; Wu et al., 2016; Yan et al., 2016). This approach allows visualization of individual reporter mRNAs via MS2 stem-loops and simultaneous visualization of translation via the SunTag, which consists of an array of antibody epitopes that are recognized by genetically-encoded single-chain antibodies (scFv). To accomplish this, we engineered a HeLa cell line stably expressing GFP-tagged scFv and Halo-tagged MS2 coat protein (MCP) (Voigt et al., 2017). The reporter mRNA encodes the SunTag array in frame with *Renilla* luciferase, enabling the measurement of protein synthesis by two complementary assays. The reporter mRNA further contains a destabilized FKBP domain to enhance the degradation of the mature protein, thereby facilitating imaging of nascent peptides during translation. To enable visualization of translation during stress, we further placed the 5’UTR of ATF4 in front of the SunTag array (Figure 1A). This enables the mRNA to be efficiently translated during stress conditions when many other mRNAs are inhibited. The reporter mRNA was integrated into the genome at a site-specific locus and was under the control of a doxycycline-inducible promoter, which enables precise mRNA tracking and minimizes potential artifacts of overexpression. Using luciferase assays, we confirmed that the translation of this ATF4-SunTag reporter is specifically upregulated during arsenite stress and repressed under normal growth conditions (Figure 1B), as previously reported for endogenous ATF4 (Andreev et al., 2015; Roybal et al., 2005). Importantly, ATF4-SunTag was efficiently translated at an oxidative stress condition (100 μM arsenite) that also triggered robust SG formation in all cells (Figure 1B), enabling us to visualize the translation of this mRNA in SG-containing cells. It should be noted that at higher arsenite concentrations, ATF4-SunTag translated was inhibited.

**Figure 1.**
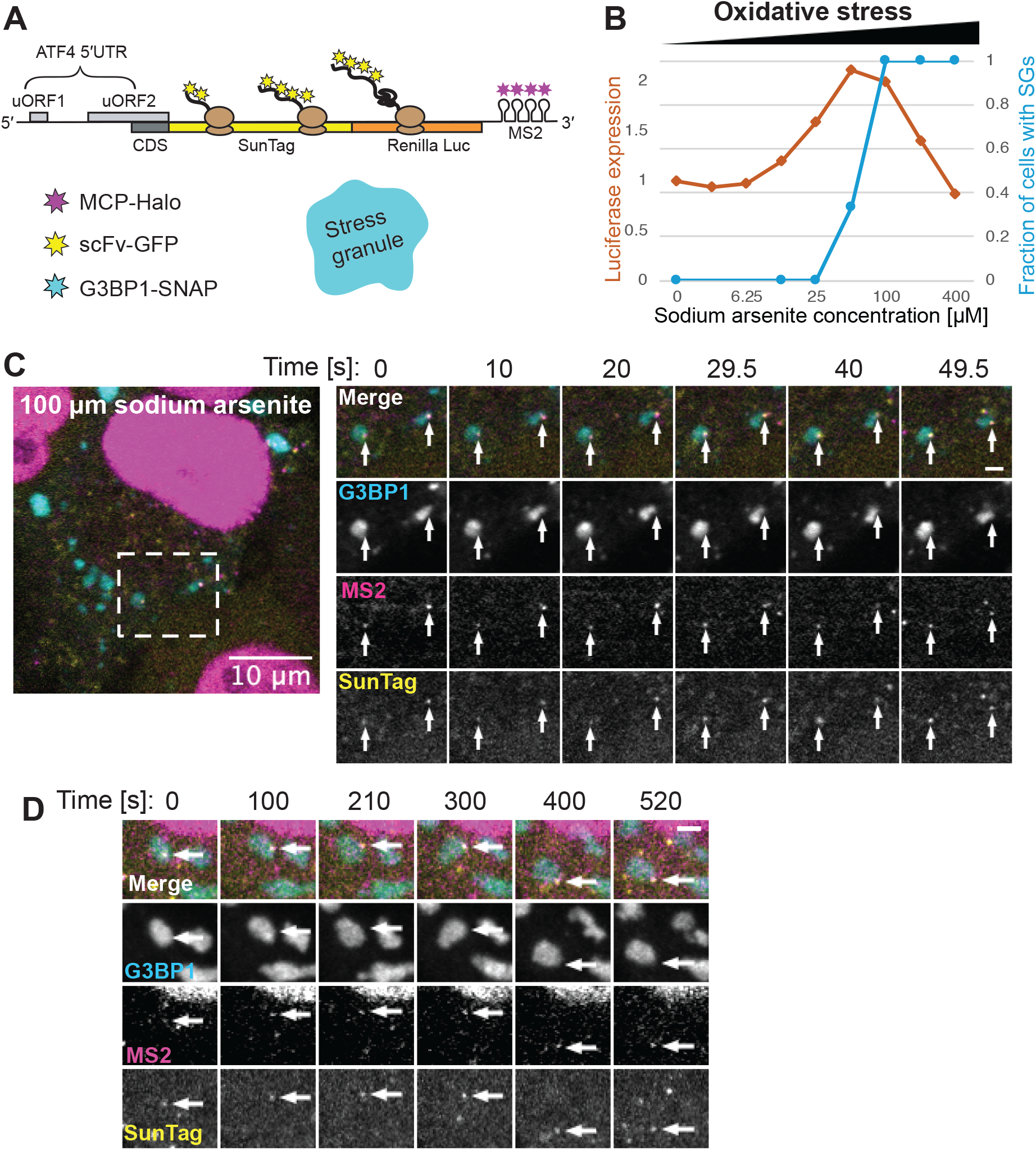
Single-molecule imaging reveals mRNA translation associated with SGs. (A) Schematic representation of ATF4-SunTag reporter mRNA used for single-molecule imaging of mRNA translation during stress. The construct is integrated in HeLa cells stably expressing MCP-Halo, scFv-GFP, and G3BP1-SNAP. (B) HeLa cells carrying the ATF4-SunTag reporter, MCP-Halo and scFv-GFP were treated with different concentrations of sodium arsenite for 1 h, and expression of the reporter was measured with luciferase assays. Fractions of cells with SGs were determined by immunofluorescence staining of G3BP1 in the same cell line exposed to the same conditions. (C) Example of SG-associated translation in cells treated with 100 μM sodium arsenite for 1 h. Excerpts from a time-lapse movie are shown, with arrows indicating SunTag-positive ATF4-SunTag mRNAs, and scale bar indicating 2 μm. (D) Excerpts from a time-lapse movie with 10 s intervals and Z-stack acquisition, showing long-lasting localization of SunTag-positive mRNA to SG (arrows). Cells were treated with 100 μM sodium arsenite for 1 h. Scale bar = 2 μm.

Since ATF4 translation is enhanced during stress, it is interesting that it has been previously reported that the chemotherapeutic drug sorafenib, which triggers the stress response, results in accumulation of ATF4 mRNA in SGs (Adjibade et al., 2015). This paradoxical finding was interpreted as a means to buffer ATF4 translation during stress by sequestration and translational repression in SGs. Using single-molecule FISH combined with immunofluorescence, we confirmed that endogenous ATF4 mRNAs and the ATF4-Suntag transcripts accumulate in SGs also during oxidative stress (Figure S1A and S1B). Next, to label SGs in living cells, we introduced SNAP-tag-labelled G3BP1 to our cell line and confirmed that G3BP1-SNAP forms SGs only during stress and colocalizes with other SG components such as endogenous TIA-1 (Figure S1C). We then performed live-cell imaging of this cell line exposed to oxidative stress and observed that ATF4-SunTag mRNAs are frequently recruited to SGs and stably associate as described for other mRNAs that are translationally inhibited (Moon et al., 2019; Wilbertz et al., 2019). SG-localized mRNAs can be clearly distinguished from cytosolic mRNAs, as they colocalize with G3BP1-labelled SGs and are much less mobile compared to freely-moving cytosolic mRNAs (Video S1). Translating and non-translating mRNAs can be distinguished by the presence of the SunTag signal. As expected, we often observe non-translating mRNA localized to SGs (Figure S1D).

Surprisingly, we also observe that translating mRNAs are frequently localized to SGs (Figure 1C, Video S1), suggesting that localization to SGs does not require inhibition of mRNA translation. Because SG-localized mRNAs are relatively static, we were able to follow them in longer time-lapse imaging experiments that revealed translating mRNAs often remain localized to SGs for more than five minutes (Figure 1D, Video S2).

### SG-associated translation is not a rare event

Considering that we observe both non-translating and translating mRNAs in SGs as well as in the cytosol, we set out to quantify whether there is any correlation between translational status and localization to SGs. To this end, we developed an image analysis workflow to perform single-particle tracking and colocalization analysis that enabled us to assign translational status and localization to thousands of ATF4-SunTag mRNAs. This analysis revealed substantial cell-to-cell variability in the fraction of mRNAs undergoing translation. On average, 30% of SG-localized ATF4-SunTag mRNAs were found to co-localize with SunTag signal, demonstrating that SG-associated translation is not a rare event (Figure 2A). Cytosolic mRNAs had a slightly higher percentage of translating mRNAs (44% of mRNAs). After addition of the translational inhibitor puromycin, almost all mRNAs (both cytosolic and SG-localized) are devoid of SunTag signal, demonstrating that our analysis is stringent and SunTag-labelled mRNAs represent *bona fide* translation sites (Figure 2A). Importantly, SG-localized and cytosolic mRNAs did not dramatically differ in their SunTag signal intensities (Figure 2B). As SunTag intensity correlates with the number of ribosomes, this indicates that ribosome occupancy on translating mRNAs is similar in the cytosol and SGs.

**Figure 2.**
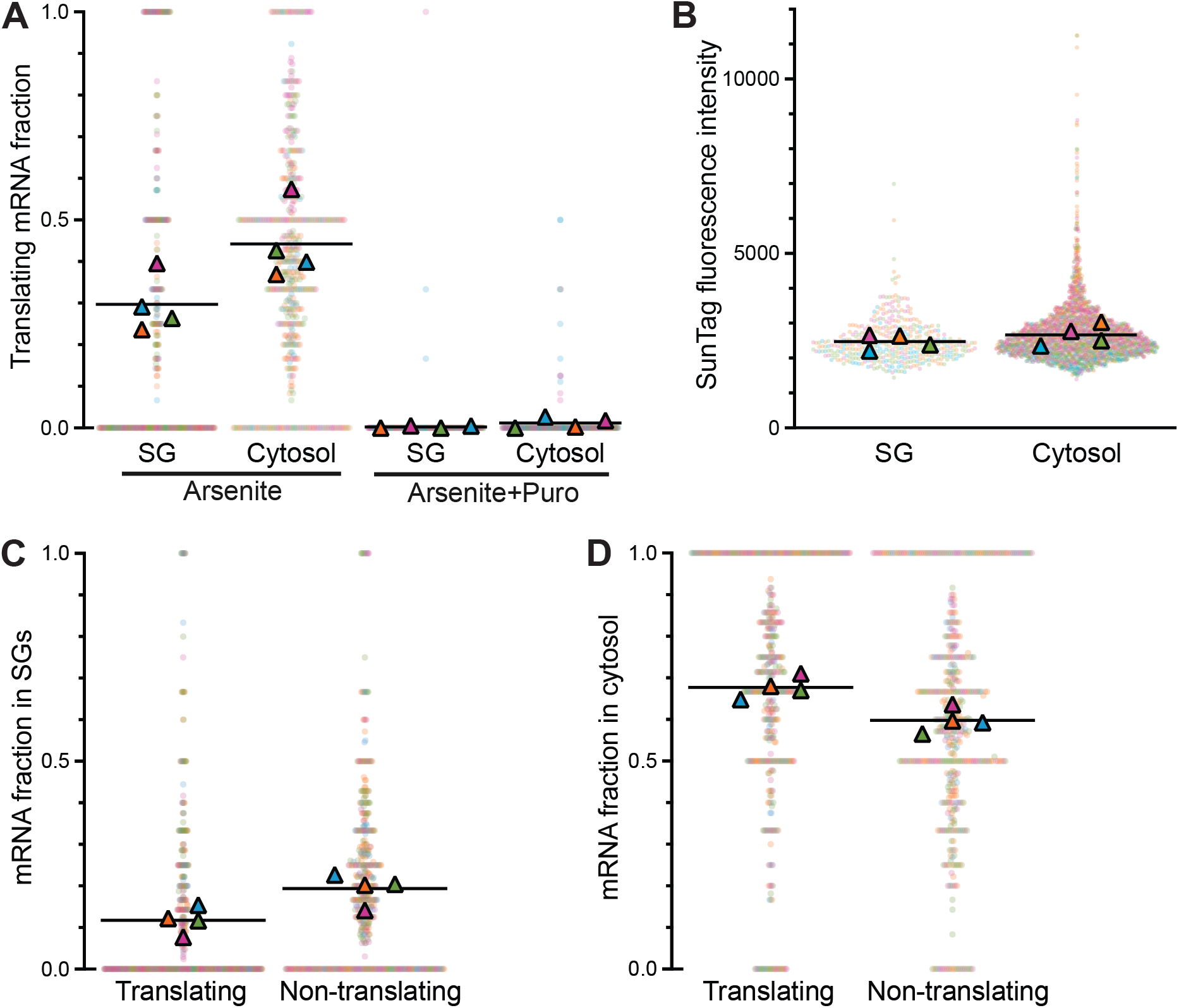
Quantification of mRNA localization and translation based on single-particle tracking. (A) Fraction of ATF4-SunTag mRNAs undergoing translation was quantified separately for mRNAs localized to SGs or mRNAs localized to cytosol. Dots represent fractions in individual cells, each color-coded by experiment. Triangles represent global fractions per experiment, and black line indicates their mean value. Cells were treated with 100 μM sodium arsenite for 50-70 min (Arsenite: 517 cells, 9023 mRNA tracks). 100 μg/mL puromycin was added before imaging where indicated (Arsenite+Puro: 264 cells, 3493 mRNA tracks). Mean +/− SEM (Arsenite): 30% +/− 3.5% (SG) vs 44% +/− 4.5% (Cytosol); p = 0.0027. (B) Mean fluorescence intensities of SunTag foci colocalized with SG-localized mRNAs and cytosolic mRNAs. Values are plotted for each cell (dots) as well as global fractions per experiment (triangles). Same dataset as in A (Arsenite). p = 0.0611. (C) Fraction of mRNAs localized to SGs was quantified separately for translating mRNAs and non-translating mRNAs. Same dataset as in A (Arsenite). Mean +/− SEM: 12% +/− 1.6% vs 19% +/− 1.8%; p = 0.0006. (D) Fraction of mRNAs localized to cytosol was quantified separately for translating mRNAs and non-translating mRNAs. Same dataset as in A (Arsenite). Mean +/− SEM: 68% +/− 1.3% vs 60% +/− 1.5%; p = 0.0045. The sum of fractions in C and D is less than 1, as some mRNA tracks were not assigned localization due to ambiguity (see Methods). Fractions in cells can cluster at discrete values (e.g., 0, 0.5, 1) due to low number of mRNAs in some cells.

To further examine how mRNAs are distributed based on their translational status, we divided all ATF4-SunTag mRNAs into translating and non-translating populations, and quantified their distribution in SGs and the cytosol. For both translating and non-translating transcripts, only a small percentage of these two populations were localized to SGs (translating: 12%, non-translating: 19%) while the majority of both types of transcripts remained in the cytosol (Figure 2C and 2D). The slight enrichment of non-translating mRNAs in SGs can have two possible explanations. One possibility is that the localization to SGs reduces the translation activity of mRNAs. The second explanation is that the ribosome occupancy of mRNAs influences the partitioning between cytosol and SG. To test this, we compared the extent of mRNA localization to SGs between arsenite-treated cells and cells cotreated with puromycin. This analysis revealed that puromycin-treated cells have a significantly higher fraction of SG-localized mRNAs (36% vs 16%), indicating that nontranslating mRNAs are more likely to be recruited to SGs (Figure S2). This strongly suggests that the observed correlation between localization to SGs and slightly lower translation activity can be explained by the higher propensity of non-translating mRNAs to localize to SGs. As the majority of mRNAs are translationally repressed during stress, this explains why previous experiments were unable to separate SG localization from translation inhibition.

### Position of mRNAs within SGs is similar for translating and repressed transcripts

It has been reported that SGs are not homogenous and contain substructures. In particular, several studies support a model where SGs contain stable cores surrounded by a more dynamic shell (Cirillo et al., 2020; Jain et al., 2016; Niewidok et al., 2018). Previously, it was also shown in Drosophila embryos that *gurken* transcripts are translated on the outside edge of P-bodies, another type of membrane-less organelle, and not in their core (Davidson et al., 2016). Therefore we set out to determine if the translational status of mRNAs was related to their position within a SG. To this aim we performed a 3D spot detection and SG segmentation in fixed cells and measured the distances from each ATF4-SunTag mRNA in SGs to the closest SG boundary in 3D. It should be noted that SG boundaries are not precisely defined and it has been shown that some SG components, such as eIF4A, can extend beyond the boundary defined by G3BP1 (Moon et al., 2019; Tauber et al., 2020). Therefore our measurement of the distance from the G3BP1 boundary could be an underestimate of the position within a SG. This analysis revealed that the majority of translating and non-translating mRNAs in SGs are located within 300 nm of the detected SG boundary (79% and 66% respectively) (Figure 3A). However, some translating mRNAs were also located deeper inside SGs, up to 750 nm from the boundary in larger SGs (Figure 3B). Importantly, the distance from the SG boundary did not correlate with SunTag intensity (Figure 3C). This indicates that translational status is not a major determinant of the position of mRNAs within SGs.

**Figure 3.**
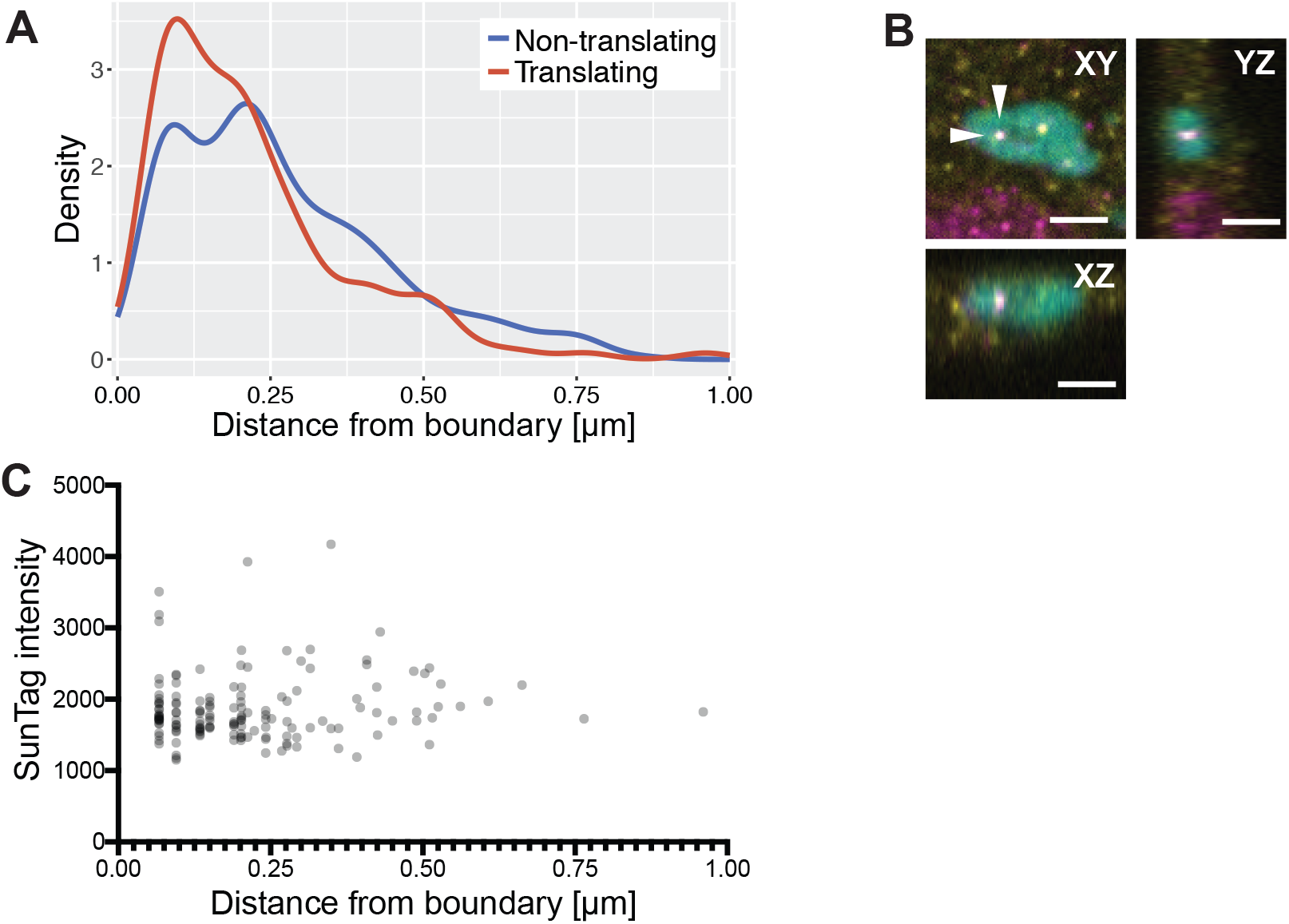
Position of mRNAs within SGs. (A) Density plot showing the distribution of distances from SG boundary for non-translating or translating ATF4-SunTag mRNAs detected within SGs. Values do not reach 0 because the measurements are pixel-based, and the smallest possible distance corresponds to the pixel size. (B) Orthogonal view of a translating mRNA in a SG. Scale bar indicates 2 μm. Merged image composed of G3BP1-SNAP (cyan), MS2-Halo (magenta), SunTag (yellow). (C) Scatter plot showing all translating mRNAs detected within SGs. Mean SunTag intensity is plotted against the distance from SG boundary.

### SG-localized mRNAs are actively translated

While our experiments demonstrate that mRNAs that are bound by ribosomes with nascent polypeptides can be localized to SGs, it remained a possibility that these transcripts were translationally stalled (Moon et al., 2020). In order to determine if SG-localized mRNAs are actively translated, we analyzed our images to identify transcripts where the SunTag signal increased over time on SG-localized mRNAs that were initially devoid of SunTag signal (Figure 4A, Video S3). These transcripts indicated that mRNAs can also initiate translation when localized to SGs.

**Figure 4.**
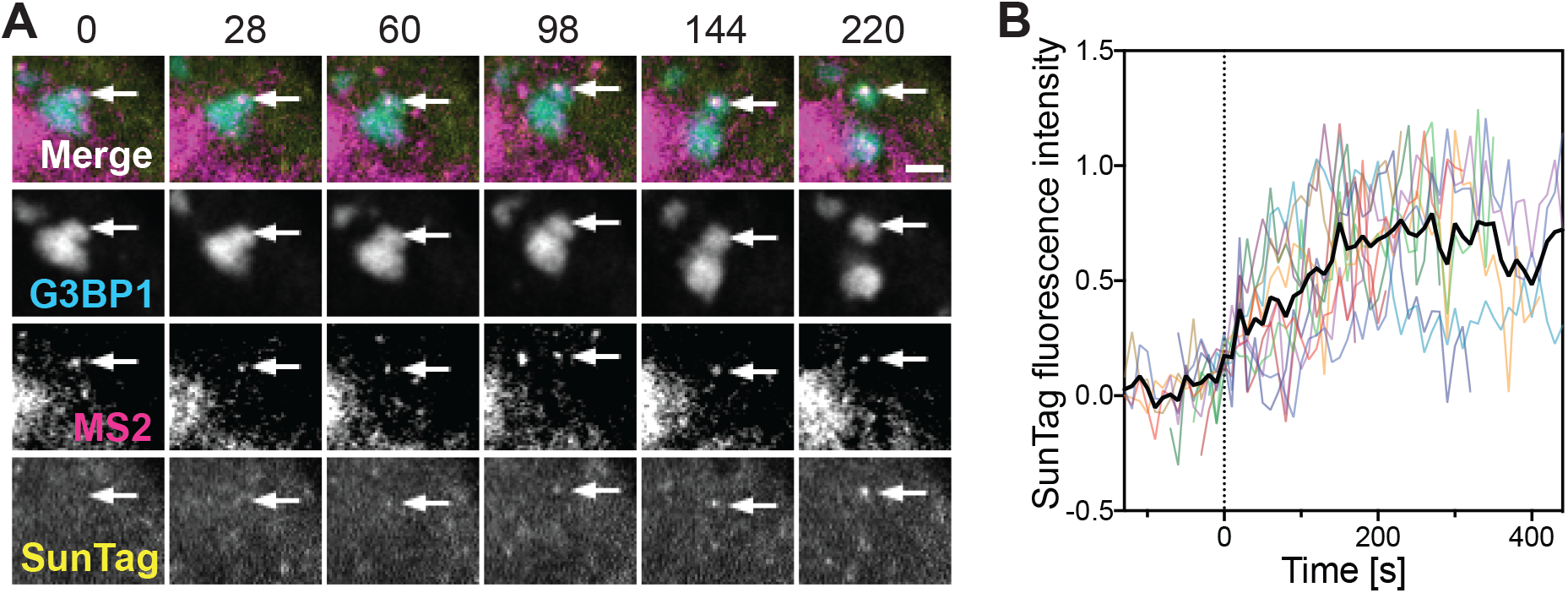
Translation dynamics on SG-localized mRNAs. (A) Excerpts from a time-lapse movie showing the buildup of SunTag signal on a SG-localized ATF4-SunTag mRNA (arrows). Cells were treated with 100 μM sodium arsenite for 1 h. Scale bar indicates 2 μm and time is shown in s. (B) Quantification of SunTag signal buildup on 11 mRNAs imaged with 10 s intervals and Z- stack acquisition. Black line indicates mean values. Curves are aligned at 0 based on the increase in SunTag signal above background levels (see Methods).

In order to quantify the kinetics of translation of transcripts localized to SGs, we acquired longer time-lapse movies of SG-localized mRNA switching from an inhibited to translating state, and we measured the increase in SunTag signal intensity (Figure 4B, S3A). The increase was not due to movement of the mRNA into the focal plane, as the MCP-Halo signal did not increase over time (Figure S3B). This quantification revealed the increase in SunTag intensity was followed by a plateau phase (Figure 4B), as previously described and can be explained by a steady state where the newly synthesized SunTag is compensated by ribosomes terminating translation (Ruijtenberg et al., 2019). The start of the plateau phase marks the first ribosome terminating translation, and measuring the duration of the increasing phase allows us to estimate the rate of translation elongation on SG-localized mRNAs. Fitting the SunTag fluorescence intensity curves with a three-part (constant-linear-constant) model determined the duration of the increasing phase to be 173 +/− 32 s (mean +/− SEM), which corresponds to an elongation rate of 5.5 +/− 1.0 codons/s. This is comparable to the elongation rates 3-5 codons/s measured by single-molecule imaging of SunTag mRNA reporters (Wang et al., 2016; Wu et al., 2016; Yan et al., 2016) or 5.6 codons/s measured by ribosome profiling (Ingolia et al., 2011), all of which were performed in the absence of stress.

To further confirm that active protein synthesis occurs on SG-localized mRNAs, we imaged the SunTag-containing SG-localized mRNAs directly after treatment with puromycin, a translational inhibitor that causes premature chain termination by incorporating into the growing peptide chain. We observed that SG-localized mRNAs lose the SunTag signal within 1 minute of puromycin addition (Figure S3C, Video S4), which is similar to previous translation site imaging experiments with membrane-anchored or cytosolic reporter mRNAs (Yan et al., 2016; Zhao et al., 2019). Together this data provides evidence that SG-localized SunTag-containing mRNAs represent sites of active translation, rather than stalled elongation complexes. Moreover, these results exclude the possibility that the interaction of translating mRNAs with SGs is mediated by the SunTag-containing peptides, as we observe the mRNA localized to SGs before the SunTag signal appears as well as after the SunTag signal disappears.

### Translating mRNAs transit between cytosol and SGs

In some of our movies, we observed the ATF4-SunTag mRNAs transitioning between the cytosol and SGs. In particular, we observed translating mRNAs moving from cytosol to SGs, where they remained localized to SGs and labelled with the SunTag signal (Figure 5A, Video S5). SG-localized translating mRNA were also observed to be released from SGs while still containing the SunTag signal (Figure 5B, Video S6). Similarly, non-translating mRNAs were also observed to transition from cytosol to SGs and from SGs to cytosol without a change in the translational status (Figure S4A and S4B). These events were relatively rare, as most mRNAs remained in either the cytosol or a SG in our movies.

**Figure 5.**
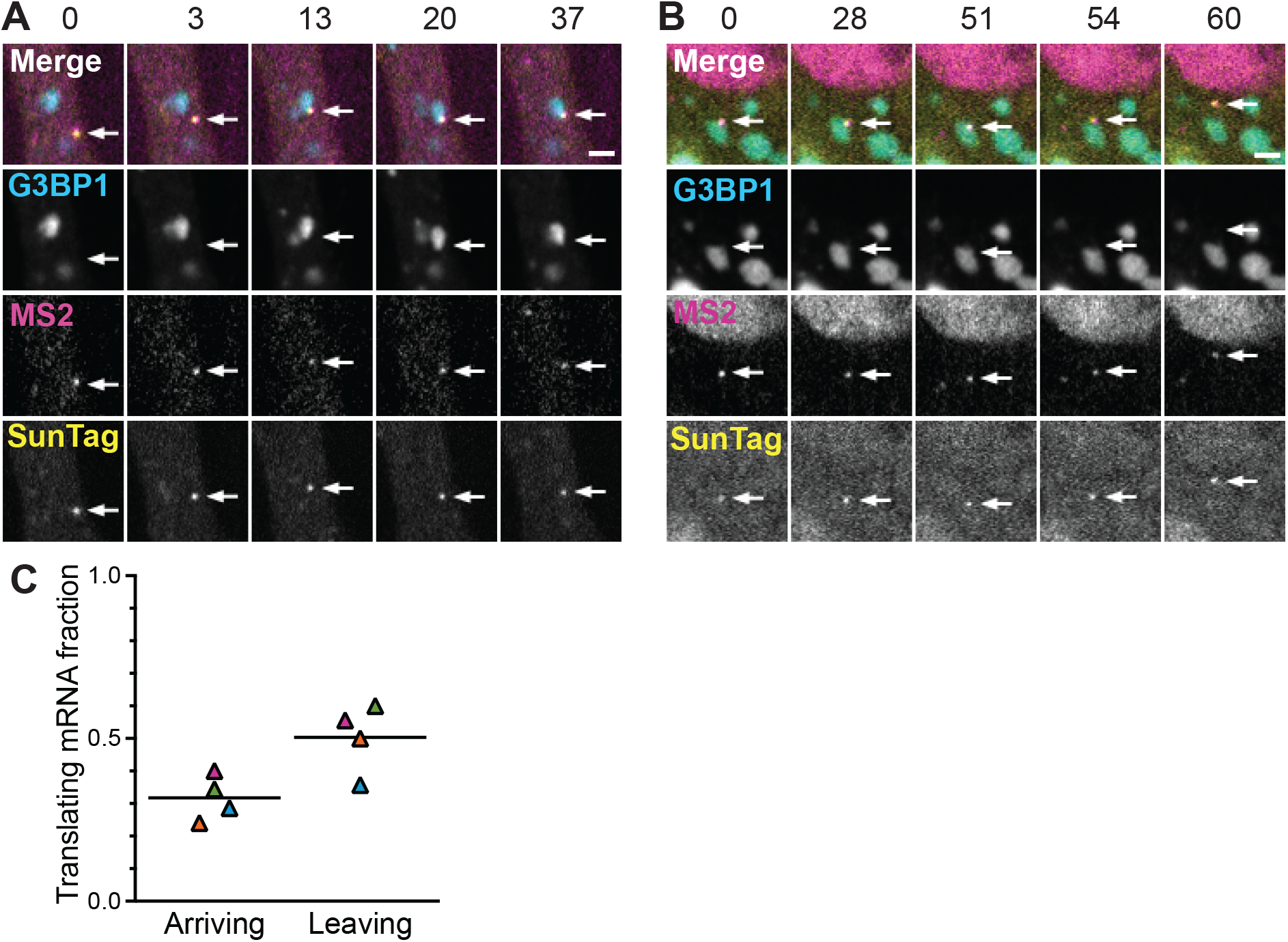
Translating mRNAs can transition between SG and cytosolic localization. (A) Excerpts from a time-lapse movie showing a translating ATF4-SunTag mRNA (arrows) arriving from cytosol to a SG. Cells were treated with 100 μM sodium arsenite for 1 h. Scale bar indicates 2 μm and time is shown in s. (B) Excerpts from a time-lapse movie showing a translating ATF4-SunTag mRNA (arrows) leaving a SG after at least 50 s localization to the SG. Cells were treated with 100 μM sodium arsenite for 1 h. Scale bar indicates 2 μm and time is shown in s. (C) Fraction of ATF4-SunTag mRNAs undergoing translation was quantified separately for mRNAs detected as arriving to SGs and mRNAs detected as leaving SGs. Fractions per experiment are plotted, with a black line representing the mean value. Same imaging dataset as in Figure 2. Mean +/− SEM: 32% +/− 3.5% vs 50% +/− 5.3%; p = 0.026.

To get more insight into the correlation between translational status and mRNA partitioning, we expanded our image analysis workflow in order to identify tracks that show clear transition between SG and cytosol within our imaging dataset. This analysis revealed that mRNAs leaving SGs are slightly more likely to be translationally active (50% of mRNA) compared to mRNAs arriving to SGs (32% of mRNA) (Figure 5C). This supports the observations that non-translating mRNAs have a higher tendency to localize to SGs (Figure S2), and that a fraction of SG-localized mRNAs can switch from non-translating to translating state before leaving SGs (Figure 4). It is important to note, however, that initiation of translation of a SG-localized mRNA does not necessarily drive its release from the SG.

### SG-associated translation is not limited to uORF-containing mRNAs

Our data indicate that SG-associated translation is relatively common for the ATF4-SunTag mRNA, which is translationally upregulated during stress due to its complex 5’UTR with two upstream open reading frames (uORFs) (Vattem and Wek, 2004). Although such uORFs are present in ~50% of the human transcriptome (Calvo et al., 2009; Johnstone et al., 2016), many mRNAs have a simpler 5’UTR and are translationally repressed during stress. One important group are 5’TOP mRNAs, which constitute up to 20% of all mRNAs in vertebrate cells and are characterized by the presence of the 5’ terminal oligo pyrimidine (5’TOP) cis-acting element (Iadevaia et al., 2008). 5’TOP mRNAs undergo strong repression during cellular stress and concomitant accumulation in SGs (Damgaard and Lykke-Andersen, 2011; Li et al., 2018; Wilbertz et al., 2019). Furthermore, it has been shown that repression of 5’TOP mRNA translation during amino acid starvation is mediated by the SG components TIA-1 and TIAR (Damgaard and Lykke-Andersen, 2011). Therefore, 5’TOP mRNAs are a prominent example of mRNAs that are assumed to be repressed in SGs (Fonseca et al., 2018; Ivanov et al., 2011).

To determine if SG-associated translation occurs for mRNAs other than ATF4 that are also repressed during stress, we used a SunTag reporter mRNA harboring a 5’TOP motif from the RPL32 5’UTR. Similarly to the ATF4-SunTag system described above, we generated a cell line carrying the 5’TOP-SunTag reporter under doxycycline-inducible promoter and stably expressing the scFvGFP, MCP-Halo and G3BP1-SNAP (Figure 6A). Using luciferase assays, we confirmed that the translation of the 5’TOP reporter is suppressed during increased levels of oxidative stress, concomitant with SG formation (Figure 6B). Using SunTag imaging, we measured that at least 76% of 5’TOP-SunTag mRNA are undergoing active translation under normal conditions. The fraction of translating mRNA was reduced dramatically to less than 2% during arsenite stress treatment (Figure 6C). Interestingly, the analysis of translation during stress revealed only 6 translating cytosolic mRNAs (out of 292 cytosolic mRNA) and 2 translating SG-localized mRNAs (out of 169 SG-localized mRNA) (Figure 6C, 6D, Video S7). Although this dramatic inhibition of translation during stress prevented us from performing a more detailed analysis, the data demonstrate that translation of 5’TOP mRNA is similarly rare in the cytosol as in SGs. In other words, SG-associated translation of 5’TOP mRNA is rare, but not due to SG-localization *per se.* In this case, SG-associated translation is rare because the conditions that permit SG formation simultaneously lead to strong translation repression for this class of transcripts.

**Figure 6.**
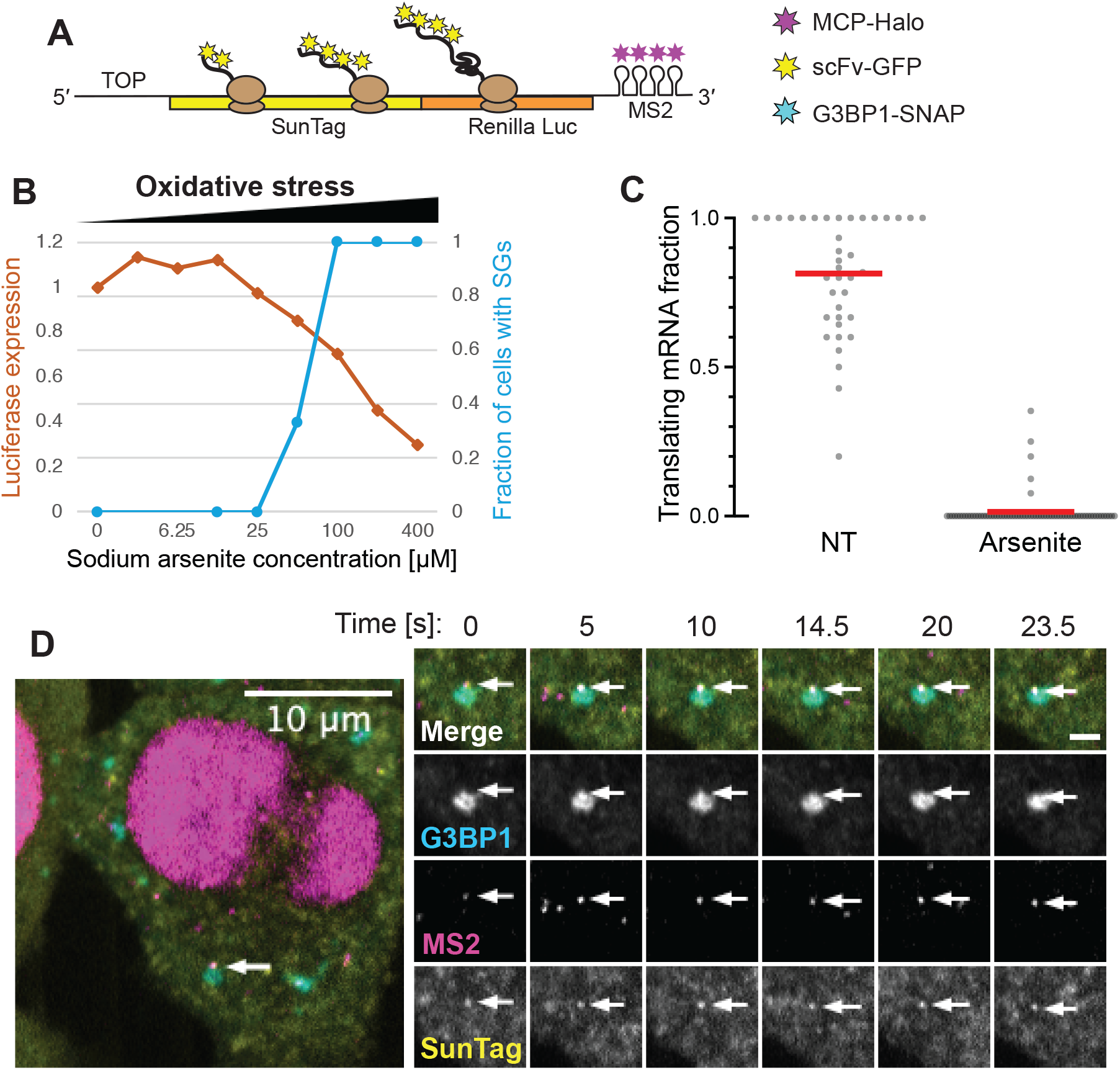
SG-associated translation of 5’TOP SunTag mRNAs during stress. (A) Schematic representation of 5’TOP-SunTag reporter mRNA used for single-molecule imaging of mRNA translation. The construct is integrated in HeLa cells stably expressing MCP-Halo, scFv-GFP, and G3BP1-SNAP. (B) HeLa cells carrying the 5’TOP-SunTag reporter, MCP-Halo and scFv-GFP were treated with different concentrations of sodium arsenite for 1 h, and expression of the reporter was measured with luciferase assays. The values are plotted together with the fractions of cells with SGs at the same conditions, adapted from Figure 1B. (C) Fraction of mRNAs undergoing translation was quantified for each cell, after 40-60 min incubation in presence (Arsenite: 70 cells, 566 mRNA tracks) or absence (NT: 38 cells, 208 mRNA tracks) of 100 μM sodium arsenite. Red line indicates mean value. (D) Example of SG-associated translation of the 5’TOP-SunTag mRNA in cells treated with 100 μM sodium arsenite for 50 min. Excerpts from a time-lapse movie are shown, with arrows indicating a SunTag-positive mRNA and scale bar indicating 2 μm.

### ATF4-SunTag mRNAs are not translated in P-bodies

P-bodies (PBs) are another type of cytoplasmic membraneless organelle believed to contain non-translating mRNAs (Teixeira et al., 2005). PBs and SGs are often located in close physical proximity and share some of their components (Kedersha et al., 2005; Matheny et al., 2019). To test whether our ATF4-SunTag reporter mRNAs are also translated in PBs, we exchanged the SG marker (G3BP1-SNAP) for a PB marker (SNAP-DCP1a) and performed similar imaging and analysis as described above. In contrast to SGs, we did not observe a stable association between the ATF4-SunTag mRNA and PBs during arsenite stress (Figure 7A and 7B). We often observe the mRNAs localized close to PBs but rarely overlapping, possibly reflecting localization to SGs (Figure 7A). We further observe mRNAs transiently touching PBs (Figure 7C, Video S8), but not forming a stable association. Although we cannot exclude that other mRNA species could be translated in PBs, this data suggests that the SG-associated translation observed for ATF4-Suntag does not occur in PBs.

**Figure 7.**
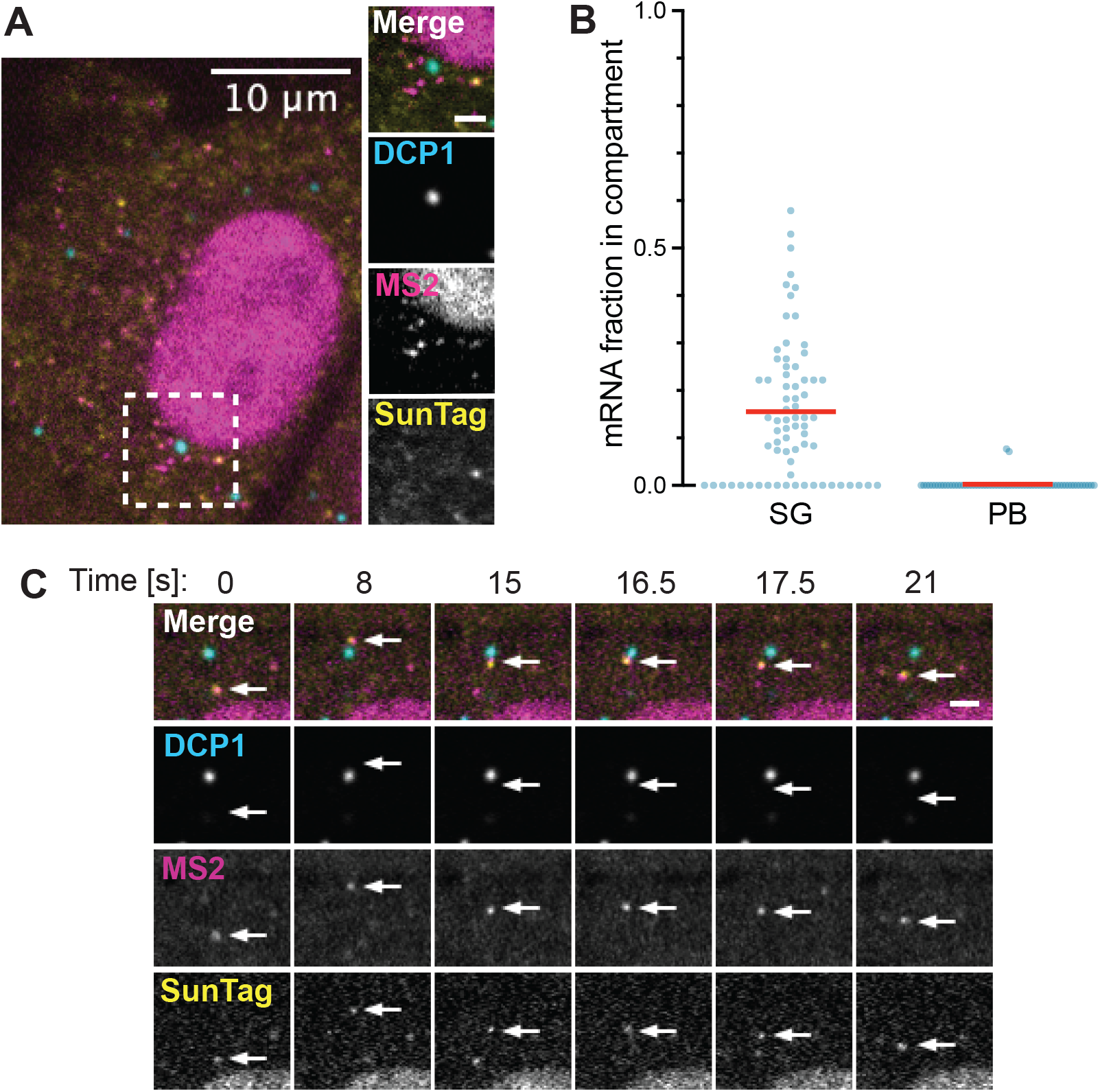
ATF4-SunTag mRNA does not show stable association with P-bodies. (A) HeLa cells carrying the ATF4-SunTag reporter together with MCP-Halo and scFv-GFP were transfected with SNAP-DCP1a to allow visualization of PBs. Live cells are imaged after 1 h treatment with 100 μM sodium arsenite. mRNAs can be seen localizing close to, but not overlapping with, PBs. 2 μm scale bar shown on right. (B) Fraction of ATF4-SunTag mRNAs localized to PBs was quantified from the sample described in A (64 cells, 881 mRNA tracks). Fraction of ATF4-SunTag mRNAs localized to SGs was quantified from cells transfected with G3BP1-SNAP instead of SNAP-DCP1a, otherwise treated the same way (71 cells, 1433 mRNA tracks). Red line indicates mean value. (C) Excerpts from a movie showing a translating ATF4-SunTag mRNA (arrows) transiently touching a PB. Same sample as described in A. Scale bar indicates 2 μm and time is shown in s.

## DISCUSSION

It is widely believed that SGs contain exclusively non-translating mRNA, leading to the assumption that all mRNAs observed in SGs (e.g. using FISH or sequencing-based methods) are in a translationally repressed state (Adjibade et al., 2015; Mollet et al., 2008; Namkoong et al., 2018). This is based on the fact that SG formation coincides with general translational inhibition during stress, and on immunofluorescence analysis of SGs that concluded that SGs are completely devoid of 60S ribosomal subunits. Antibodies against the 60S components L5, L37 and Po did not show enrichment in SGs, while antibodies against 40S components showed strong signal in SGs (Kedersha et al., 2002; Kimball et al., 2003). Consequently, the current model postulates that SG-localized mRNA are bound only by 40S subunit, representing “stalled preinitiation complexes” (Anderson and Kedersha, 2008; Ivanov et al., 2019; Panas et al., 2016; Wolozin and Ivanov, 2019). We believe this model holds true for most SG-localized mRNA, however, our data indicate it cannot be generalized for all mRNA localized to SGs and that mRNA localization to SGs does not require or cause translation inhibition.

Interestingly, there is evidence in the literature for the presence of 60S subunits in SGs. First, the immunofluorescence stainings for L5 and L37 proteins showed signal overlapping with SGs, albeit weaker than the signal of 40S components (Kimball et al., 2003). Second, a recent study has shown that 60S subunits can also accumulate in SGs during certain conditions that impair the protein quality control machinery (Seguin et al., 2014). We propose that most SG-localized mRNAs are indeed “stalled preinitiation complexes”, while some can recruit 60S subunits and undergo translation.

Importantly, the data presented here demonstrate that localization to SGs does not prevent mRNA translation. Evidence is now emerging that SGs do not have a large effect on global translation. Transcriptomic studies revealed that only ~10% of mRNA molecules accumulate in SGs (Khong et al., 2017) and it has been shown that G3BP-deficient cells are able to inhibit bulk translation during stress without forming SGs (Kedersha et al., 2016). Our study indicates that even the mRNAs recruited to SGs are not translationally repressed by the SG environment. Although we see a slight over-representation of non-translating mRNAs in SGs, this can be explained by the preferential recruitment of non-translating mRNAs to SGs (Figure S2), rather than silencing of translation in SGs. This observation of preferential recruitment of non-translating mRNAs is consistent with previous reports. For example, mRNA accumulation in SGs has been shown to correlate with the expected time of ribosome runoff from mRNAs of different lengths (Khong and Parker, 2018; Padrón et al., 2019). Importantly, our analysis of single mRNA translation dynamics indicates that localization to SGs does not prevent translation initiation and does not prevent efficient elongation. Taken together the data argue that SGs have a preference for recruitment of inhibit mRNAs, but SGs do not directly repress mRNA translation.

Membrane-less organelles or biomolecular condensates are now thought to partition cellular environments in order to enable a wide variety of biological processes to be specifically regulated (Banani et al., 2017; Boeynaems et al., 2018). This study demonstrates that it is necessary to quantitatively measure the biological processes both inside and outside of the organelle in order to understand its function. The continued development of single-molecule imaging techniques and fluorescent biosensors coupled with the identification of separation-of-function mutants or chemical biology tools will play a key role towards understanding the function of biomolecular condensates.

## ACKNOWLEDGEMENTS

This work was supported by the Novartis Research Foundation (J.A.C), the Swiss National Science Foundation grant 31003A_182314 (J.A.C), the SNF-NCCR RNA & Disease (J.A.C) and an EMBO postdoctoral fellowship (D.M). The authors thank members of the Chao lab for helpful discussions and critical reading of the manuscript, L. Gelman and S. Bourke for microscopy support, and H. Kohler for cell sorting. The authors further thank members of the S. Alberti lab and D. Schübeler for helpful feedback on the manuscript.

## AUTHOR CONTRIBUTIONS

D.M. performed all experiments and image analysis. B.E. generated the G3BP1-SNAP virus. J.E. helped with image analysis workflows. G.R. helped with calculating elongation rates. J.A.C. conceived the project. D.M. and J.A.C. wrote the article with input from all the authors.

## MATERIALS AND METHODS

### Cell culture

HeLa cells used in this study are derived from the previously-described HeLa-11ht cell line (Weidenfeld et al., 2009), which contains a site for Flp-RMCE (recombinase-mediated cassette exchange), allowing a single-copy genomic integration of a target gene. The cell line is further expressing a reverse tetracycline controlled transactivator (rtTA2S-M2), allowing doxycycline-inducible expression of the inserted gene. HeLa cells were cultured in Dulbecco’s Modified Eagle Medium (DMEM) containing 4.5 g/L glucose, Penicillin (100 U/ml),

Streptomycin (100μg/ml), L-Glutamine (4mM) and 10% fetal bovine serum (FBS). Cells were maintained at 37°C and 5% CO_2_. For transient transfections, we used Lipofectamine 2000 transfection reagent (Invitrogen) with Opti-MEM reduced serum medium (Gibco) according to manufacturer’s instructions, but scaled down to 250 ng DNA and 1 μL Lipofectamine per 1 mL growth media.

### DNA constructs

To generate the ATF4-SunTag reporter, we have modified the previously-described SunTag-*Renilla-MS2* plasmid (Wilbertz et al., 2019) (Addgene plasmid #119945) by inserting the cDNA of human ATF4 5’UTR (until the end of uORF2, thereby including the first 27 amino acids of the main coding sequence that overlap with uORF2) using HindIII and RsrII restriction sites. The resulting plasmid contains a Tet-CMV (cytomegalovirus) promoter followed by ATF4 5’UTR, SunTag(24xGCN4), *Renilla* luciferase, destabilized FKBP domain (Banaszynski et al., 2006) with a stop codon, 24xMS2 stem-loop cassette, CTE (constitutive transport element) and SV40 poly(A) tail. The main coding sequence of ATF4 is in frame with SunTag and *Renilla* luciferase. For detection of SGs and PBs, we used the previously-described lentiviral vectors carrying G3BP1-SNAP (Addgene plasmid #119949) (Wilbertz et al., 2019) and SNAP-DCP1a (Horvathova et al., 2017), respectively.

### Cell line generation

The ATF4-SunTag reporter was stably integrated into a previously-described HeLa-11ht cell line stably expressing GFP-tagged single-chain antibodies (scFvGFP) against GCN4 (Addgene plasmid #104998) and NLS-stdMCP-stdHalo fusion proteins (Addgene plasmid #104999) (Voigt et al., 2017). The genomic integration was achieved via RMCE as previously described (Weidenfeld et al., 2009). In brief, 3 x 10^5^ HeLa cells were seeded into a 6-well plate and next day transfected with 2 μg of the FLPe recombinase plasmid (Addgene #20733) (Beard et al., 2006) together with 2 μg of the plasmid carrying the SunTag reporter flanked by Flp-recombinase target sites. The next day, 5 μg/mL puromycin (Invivogen) was added to select for transfected cells. Two days later, the puromycin-containing medium was removed and cells were kept in growth medium containing 50 μM ganciclovir (Sigma-Aldrich) for 10 days to select for cells with successful RMCE. In order to generate a single-clone cell line, single-cell sorting into a 96-well plate was performed. The cells were then expanded and reporter expression was tested with luciferase assays. The cell line carrying the 5’TOP element-containing SunTag reporter (Addgene plasmid #119946) together with scFv-GFP and NLS-stdMCP-stdHalo was described previously (Wilbertz et al., 2019). G3BP1-SNAP was then stably integrated into reporter-expressing cell lines (ATF4-SunTag or 5’TOP-SunTag) using a third-generation lentiviral system (Mostoslavsky et al., 2006). Lentivirus production and infection was performed under BL-2 (biosafety level 2) conditions. Lentivirus production was performed by transfecting HEK293T cells with the G3BP1-SNAP in pHAGE lentiviral backbone along with the accessory plasmids VSV-G, Gag/Pol, Tat, Rev, using Fugene 6 (Promega). Supernatant was harvested from the transfected cells for the next three days and then centrifuged, filtered through 0.45 μM filter (Millipore) and concentrated using LentiX concentrator (Clontech). HeLa cells were then infected with the lentivirus, incubated for 6 days, passaged and subsequently FACS-sorted to select positive cells.

### Luciferase assays

*Renilla* Luciferase Assay System (Promega) was used for measuring the production of SunTag reporters. Cells carrying the ATF4-SunTag or 5’TOP-SunTag reporters (both also expressing scFv-GFP and NLS-stdMCP-stdHalo) were seeded on 24-well plates. The next day, expression of reporters was induced with 1 μg/mL doxycycline for 2 hours. The medium was then washed out with PBS and replaced with fresh medium containing different concentrations of sodium arsenite (Sigma). After 1 h incubation, the cells were lysed with 150 μL Passive Lysis Buffer per well. Lysate (30 μL) was then transferred to wells of a 96-well EIA/RIA Plate (Costar) in triplicate. Bioluminescence measurements were performed using the Mithras LB 940 microplate reader (Berthold), injecting 12 μL of *Renilla* Luciferase Assay Reagent per well. The means are calculated based on 3 biological replicates (each consisting of 3 technical replicates).

### Immunofluorescence

Cells were plated on glass coverslips (High Precision, 170 μm thickness, 18 mm diameter, Paul Marienfeld GmbH) in 12-well plates. Two days later, cells were treated as indicated, followed by a quick wash in PBS and fixation in 4% paraformaldehyde (Electron Microscopy Sciences) in PBS for 10 minutes. Cells were then washed three times in PBS and permeabilized with 0.5% Triton X-100 diluted in PBS for 5 minutes. Afterwards, the cells were washed in PBS and incubated in 3% bovine serum albumin (BSA) (Sigma-Aldrich) diluted in PBS for 1 hour. This was followed by a 1 h incubation with primary antibodies (diluted in 1% BSA in PBS), three washes in PBS, and a 1 h incubation with secondary antibodies (diluted in 1 % BSA in PBS). The following antibodies were used: Mouse anti-G3BP1 (BD Biosciences, 611127), rabbit TIA-1 (Abcam, ab40693), rabbit DDX6 (Bethyl, A300-461A). Corresponding secondary antibodies conjugated with Alexa Fluor (488, 568 or 647) dyes (Life Technologies) were used. After incubation with secondary antibodies, the coverslips were washed three times with PBS and mounted on microscope slides (VWR International) in DAPI-containing Prolong Gold (Invitrogen). Samples were imaged on the spinning-disk confocal microscope described in the “Live cell imaging” section, using single-camera sequential acquisition. Fiji (Schindelin et al., 2012) was used for image processing (cropping, brightness adjustment, merging channels, labelling with scale bars). Fractions of cells with SGs were counted manually based on immunofluorescence staining of G3BP1 after 1 h treatment with 0, 12.5 μM, 25 μM, 50 μM, 100 μM, or 200 μM sodium arsenite (715 cells were counted in total).

### Fluorescence *in situ* hybridization (FISH)

To detect endogenous ATF4 mRNA, FISH probes targeting human ATF4 (designed using the Stellaris Probe Designer) were made by enzymatic oligonucleotide labeling (Gáspár et al., 2018), using Amino-11-ddUTP (Lumiprobe) and Atto565-NHS (ATTO-TEC). To detect the ATF4-SunTag mRNA, we used Stellaris FISH probes targeting *Renilla* luciferase and labelled with the Quasar 570 dye (Biosearch Technologies). Cells were plated on glass coverslips (High Precision, 170 μm thickness, 18 mm diameter, Paul Marienfeld GmbH) in 12-well plates. Two days later, cells were treated as indicated, followed by a quick wash in PBS and fixation in 4% paraformaldehyde (Electron Microscopy Sciences) in PBS for 10 minutes. Cells were then washed three times in PBS and permeabilized with 0.5% Triton X-100 diluted in PBS for 5 minutes. Afterwards, the cells were further washed with PBS and then washed two times with wash buffer (10% formamide (Abcam) in 2x saline sodium citrate (SSC). Coverslips were then incubated for 4 h with hybridization solution (2 x SSC, 10% formamide, 10% dextran sulfate, 0.5% BSA, 200 nM FISH probes) at 37°C in a humidified chamber. Cells were then washed twice with wash buffer for 30 min and then twice with PBS. Immunofluorescence staining was subsequently performed according to the “immunofluorescence” section (from BSA blocking to mounting). Samples were imaged on the spinning-disk confocal microscope described in the “Live cell imaging” section, using single-camera sequential acquisition.

### Live cell imaging

Cells were plated on glass-bottom 35 mm μ-Dish (ibidi GmbH) or μ-Slide (ibidi GmbH). After 2 days, the growth medium was replaced with a fresh medium containing 1 μg/mL doxycycline in order to induce the transcription of reporter mRNAs. After 70 minutes, the medium was further supplemented with JF585 HaloTag ligand and JF646 SNAP-tag ligand, obtained from L. Lavis (Janelia Research Campus) (Grimm et al., 2015, 2017), each at 100 nM final concentration. After 20 minutes of further incubation, the medium was removed and cells were washed in PBS. Cells were then kept in FluoroBrite DMEM (Life Technologies) supplemented with 10% FBS and 4 mM L-glutamine (and sodium arsenite if indicated) for further 40-60 minutes and during imaging. Cells were then imaged on an inverted microscope Ti2-E Eclipse (Nikon) equipped with CSU-W1 Confocal Scanner Unit (Yokogawa), 2 back-illuminated EMCCD cameras iXon-Ultra-888 (Andor), MS-2000 motorized stage (Applied Scientific Instrumentation) and VisiView^®^ imaging software (Visitron Systems GmbH). Illumination was achieved with 561 Cobolt Jive (Cobolt), 488 iBeam Smart, 639 iBeam Smart (Toptica Photonics) lasers and VS-Homogenizer (Visitron Systems GmbH). CFI Plan Apochromat Lambda 100X Oil/1.45 objective (Nikon) was used, resulting in pixel size of 0.13 μm. Unless otherwise stated, excitation in 3 channels was performed sequentially with 45 ms exposure times in a single plane. The 488 nm channel emission was detected on the second camera. To correct for camera misalignment, images of 0.5 μm fluorescent beads were acquired using the TetraSpeck™ Fluorescent Microspheres Size Kit (Thermo Fisher Scientific) on each imaging session. During all incubation steps as well as imaging, the cells were kept at 37°C and 5% CO_2_. To perform time-lapse imaging immediately after puromycin addition, puromycin was pre-diluted in FluoroBrite and added to the μ-Slide mounted on the microscope stage with cells in the field of view and in focus.

### Image processing of live cell data

First, images of fluorescent beads were used to measure the channel shift using the Descriptor-based registration plugin (Preibisch et al., 2010) in Fiji (Rueden et al., 2017; Schindelin et al., 2012). The affine transformation model was then re-applied to the 488 nm channel in all movies acquired on the same imaging session using a Fiji macro. To create representative movies and images, Fiji was used to create maximum intensity projections (in case of multi-plane acquisition) and correct for photobleaching using the Bleach correction plugin with the “simple ratio” correction method. Fiji was further used for cropping, brightness adjustment, merging channels, labelling images with scale bars, arrows and time, and for creating montages from movies.

### Track-based analysis of translational status and localization

mRNA tracking-based quantification of translation and SG localization was performed using a custom-built workflow in KNIME (Berthold et al., 2009; Dietz et al., 2020). First, 15-frame (~ 7 s) long movies (after registration as described above) were imported. Then, ROIs were manually annotated to include cytosol (but not nucleus) of cells with visible mRNA and SGs. Tracking was then performed in both SunTag and MS2 channels using a custom-build KNIME node enabling the software to run TrackMate (Tinevez et al., 2017) in batch mode with the Multi-channel intensity analyzer module. This node is available on KNIME Node Repository: KNIME Image Processing / ImageJ2 / FMI / Track Spots (Subpixel localization, multi-channel) (Eglinger, 2019). The tracking process involved spot detection with “Laplacian of Gaussian” detector with estimated spot radius of 200 nm and sub-pixel localization. Detection thresholds were adjusted for each experiment but kept constant within the experiment. As a particlelinking algorithm, the ‘‘Simple LAP tracker” was used with maximum linking and gap-closing distances of 1200 nm and maximum gap of 1 frame. To test whether mRNA is localized to SGs, we measured the mean intensity of G3BP1-SNAP at the coordinates of each detected spot. This intensity was then divided by the mean G3BP1-SNAP intensity of the whole annotated ROI, resulting in values of G3BP1 enrichment at the mRNA foci over the cytosol. We then defined spots with G3BP1 enrichment > 1.3 as localized to SGs. Only when every spot of the track fulfills this condition, the mRNAs are defined as SG-localized. Conversely, when every spot of a track has G3BP1 enrichment < 1.3, the mRNA is defined as cytosolic. Tracks that do not fulfill either condition are not assigned localization due to ambiguity (e.g., localization at SG boundary or only transient colocalization). Exceptionally, mRNAs were defined as “arriving to SG” when at least 2 spots (including the first) were “cytosolic” and at least 4 spots (including the last) were “localized to SGs”. mRNAs were defined as “leaving SG” when at least 4 spots (including the first) were “localized to SGs” and at least 2 spots (including the last) were “cytosolic”. To test the translational status of mRNAs, we performed pairwise colocalization of MS2 and SunTag track coordinates as previously described (Voigt et al., 2018), linking the mutual nearest neighbors between the two channels within a distance cutoff of 500 nm. The MS2 tracks are then classified as translating mRNA if they contain at least 2 colocalizing spot pairs, and MS2 tracks that contain less than 4 spots are excluded from the analysis. All other MS2 tracks are classified as non-translating. Fractions of translating mRNA were then calculated from all SG-localized or all cytosolic mRNA in each cell. Similarly, the fractions of SG-localized mRNA were calculated from all translating or all non-translating mRNAs. Global fractions per experiment were further determined from total numbers of tracks (without cell-level grouping) to avoid bias towards cells with low mRNA numbers. The global fractions per experiment were plotted (triangles) and used to calculate the average values, standard error of the mean (SEM), and p-values (paired t-test). To quantify mRNA localization to PBs, we used the same workflow as for SGs, measuring SNAP-DCP1a signal instead of G3BP1-SNAP signal. Values were plotted using GraphPad Prism.

### Measurement of SunTag intensity buildup

To measure the changes in fluorescence intensity over time, measurements were performed using Fiji on registered movies. Mean intensity of SunTag and MS2 signal on mRNA foci was measured at each time point in round ROIs with 5 pixel diameter. Local background fluorescence was measured at each time point in doughnut-shaped ROIs around the foci and subtracted from the foci intensities. For each time point (t), three-frame moving averages were then calculated from t, t+1, t+2, and used for normalization and alignment of the intensity curves. First, curves were normalized by dividing each value by the highest moving average of the entire track (y = 1 at the highest moving average of each track). Second, the curves were aligned at x = 0, defined as the first time point with moving average above 0.20. Mean values were calculated from the normalized and aligned values. The intensity curves along with the mean values were plotted in GraphPad Prism. The buildup of fluorescence intensity on newly translated mRNAs includes an “increasing phase” and a “plateau phase”, with the duration of the “increasing phase” reflecting the time it takes for the first ribosome to reach the stop codon (Ruijtenberg et al., 2019). The duration of the “increasing phase” can, therefore, be used to estimate the translation elongation rate on the SG-localized mRNAs. Our measurements additionally contain an “initial phase” with background-corrected SunTag intensities close to 0, as we detect MS2 foci before the SunTag signal appears. Some of the curves also exhibited a drop in intensity after the plateau. These frames were filtered out when the 3-frame moving average decreased below 0.5 after the maximal value. In order to calculate the duration of the “increasing phase”, we then fit the data to a three-part model. We assume that the background-corrected intensity is 0 until time t_1_, then it increases at rate β between time t_1_ and t_2_, and from time t_2_ it stays constant. Let t be the time in seconds and X(t) be the signal intensity. We assume that

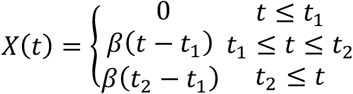

The optimal parameters β, t_1_, and t_2_ minimize the least-square error between expected and measured signal intensity. This optimization was done with the MATLAB function lsqcurvefit. To estimate the elongation rate, we then calculated the number of codons that can contribute to the increase of SunTag intensity. We counted the codons between the end of 1. GCN4 repeat and the stop codon. 30 codons were further subtracted to correct for the length of the ribosome exit tunnel (Boersma et al., 2019). The number of codons was then divided by the calculated duration of the “increasing phase” and reported in codons/s together with standard error of the mean (SEM).

### 3D distance measurements

To measure distances between mRNAs and SG boundary, cells were plated on glass coverslips (High Precision, 170 μm thickness, 18 mm diameter, Paul Marienfeld GmbH) in a 12-well plate. After 2 days, the growth medium was replaced with a fresh medium containing 1 μg/mL doxycycline in order to induce the transcription of reporter mRNAs. After 70 minutes, the medium was further supplemented with JF585 HaloTag ligand and JF646 SNAP-tag ligand, each at 100 nM final concentration. After 20 minutes of further incubation, the medium was removed, washed out with PBS, and replaced with fresh DMEM containing 100 μM sodium arsenite. After 1 h, cells were quickly washed in PBS and fixed in 4% paraformaldehyde (Electron Microscopy Sciences) diluted in PBS for 10 minutes. Cells were then washed three times in PBS and mounted on microscope slides (VWR International) in Prolong Gold (Invitrogen). Samples were imaged the next day on the spinning-disk confocal microscope described in the “Live cell imaging” section, using single-camera sequential acquisition. In addition to the CFI Plan Apochromat Lambda 100X objective, an extra 2x magnifying lens inside the scan-head in front of the camera was used, resulting in a pixel size of 67 nm. Z-stacks with 25 slices and 200 nm step size were acquired. The z-stacks were analyzed in a custom-built KNIME workflow. Briefly, G3BP1-SNAP signal was used to segment SGs in 3D using Otsu thresholding (Otsu, 1979) after applying Gaussian blur (sigmas: 2, 2, 1 pixels in X, Y, Z). Distance map was then generated, calculating the minimal distance of each foreground pixel to the nearest background pixel in 3D. MS2 foci and SunTag foci were then separately detected using a custom-built KNIME node that runs the spot detection module of TrackMate in batch mode. This node is available on KNIME Node Repository: KNIME Image Processing / ImageJ2 / FMI / Spot Detection (Subpixel localization) (Eglinger, 2019). Estimated spot radius of 200 nm was used. SunTag foci were assigned to nearby MS2 foci using recursive spot pairing as described above, in this case using 3D coordinates and a distance cutoff of 300 nm. MS2 spots with a paired SunTag spot were labelled as translating, and remaining MS2 spots were labelled as non-translating. Each mRNA (MS2 foci) was then assigned with pixel coordinates and the value on the distance map, reflecting the minimal distance to the nearest pixel outside of SG. These distance map values were then plotted for all translating and non-translating mRNAs within SGs, using the density function in R. The scatter plot of distance map values and SunTag intensities was plotted for all translating mRNAs in SGs using Prism. 3D visualizations of mRNAs in SGs were created using the Orthogonal Views function in Fiji.

---

**Figure S1.**
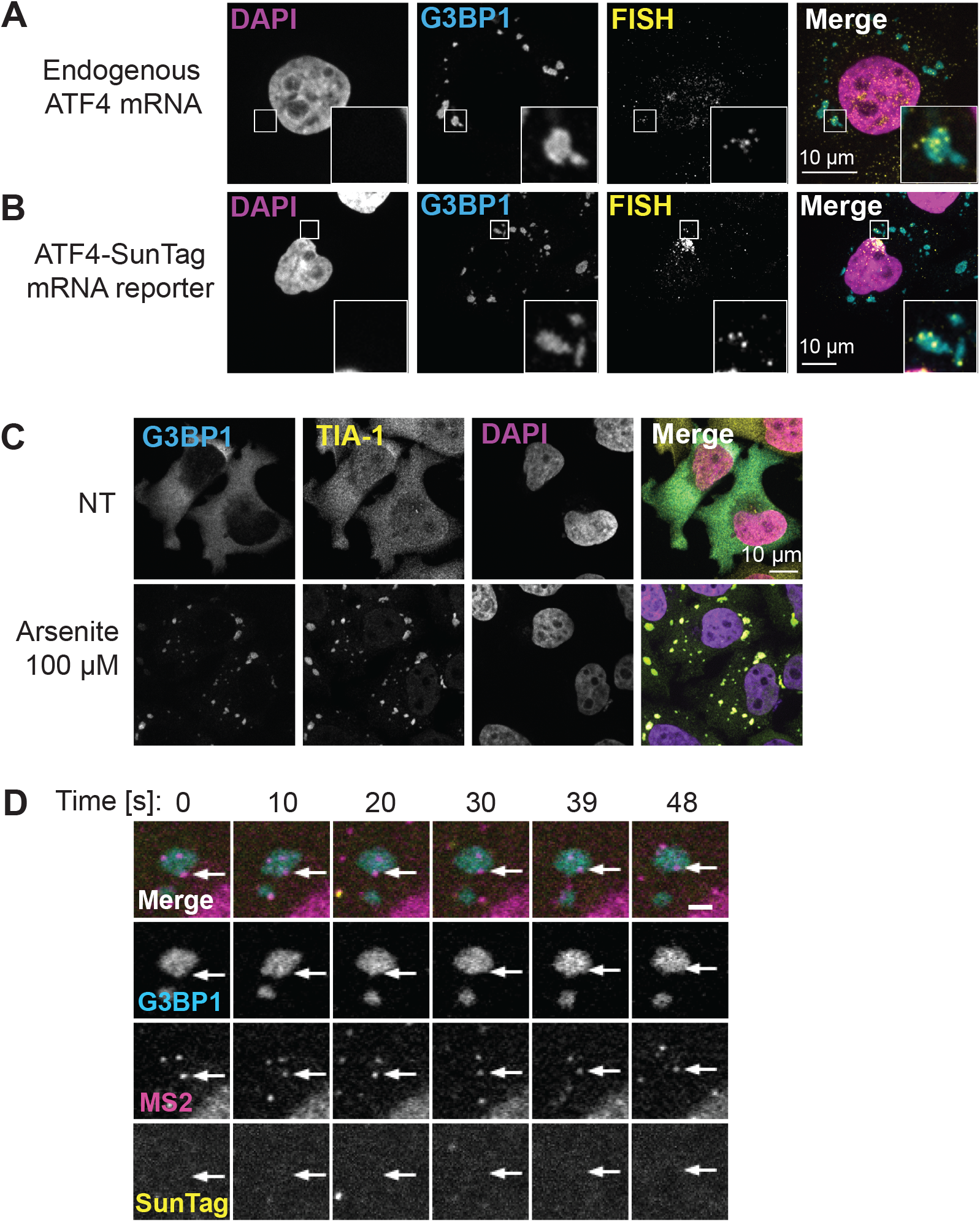
Characterization of ATF4 mRNA localization and SG formation. Related to Figure 1. (A) Endogenous ATF4 mRNA detected with FISH and G3BP1 detected with immunofluorescence in HeLa cells (parental HeLa 11ht). Cells were incubated for 1 h in the presence of 100 μM sodium arsenite. (B) ATF4-SunTag reporter mRNA detected with FISH and G3BP1 detected with immunofluorescence in HeLa cells (HeLa 11ht with stably integrated ATF4-SunTag reporter and no additional transgenes). Cells were incubated for 1 h in the presence of 100 μM sodium arsenite. (C) HeLa cells expressing G3BP1-SNAP were incubated for 1 h in presence (Arsenite) or absence (NT) of sodium arsenite. TIA-1 was detected via immunofluorescence and G3BP1 via the SNAP dye. (D) Example of non-translating mRNAs (arrows) localized to SGs in cells expressing ATF4-SunTag, MCP-Halo, scFv-GFP and G3BP1-SNAP, treated with 100 μM sodium arsenite for 1 h. Excerpts from a time-lapse movie are shown, with a scale bar indicating 2 μm.

**Figure S2.**
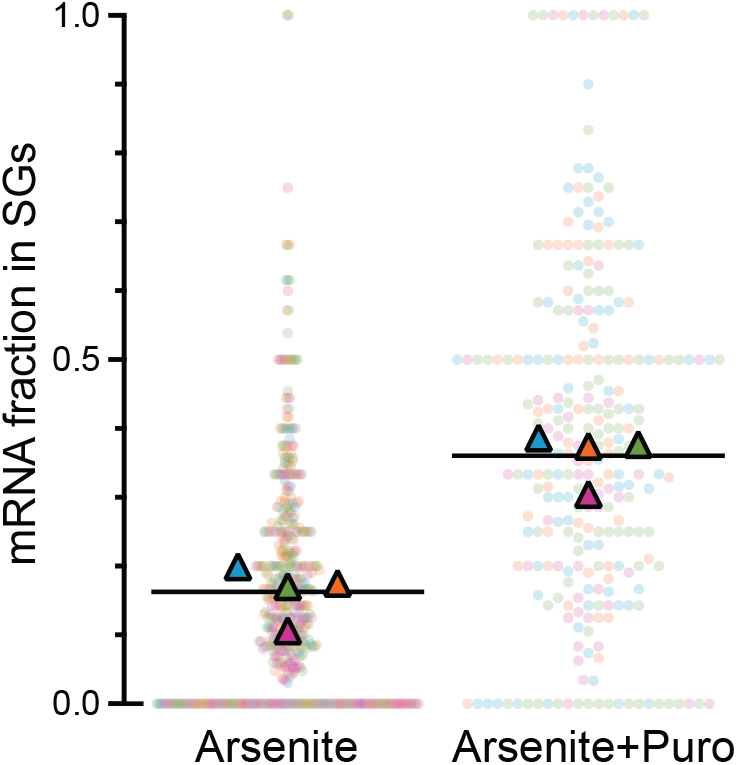
Quantification of mRNA localization to SGs. Related to Figure 2. Fraction of ATF4-SunTag mRNAs localized to SGs was measured using single-particle tracking in cells treated with 100 μM sodium arsenite for 50-70 min. 100 μg/mL puromycin (Puro) was added before imaging where indicated. Dots represent fractions in individual cells, each color-coded by experiment. Triangles represent global fractions per experiment, and black line indicates their mean value. Same dataset as in Figure 2A. Mean +/− SEM: 16% +/2.0% vs 36% +/− 1.9%; p < 0.0001.

**Figure S3.**
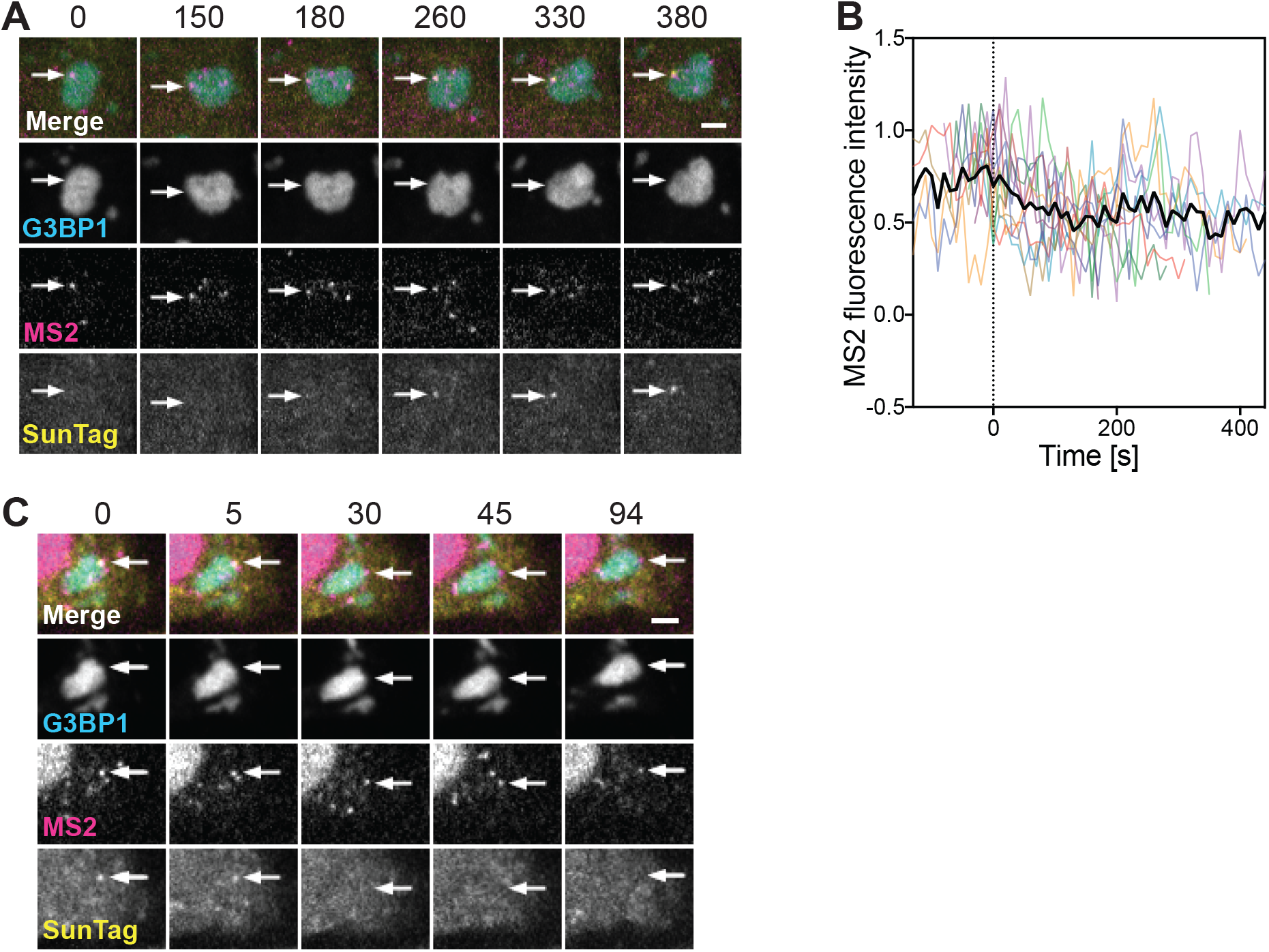
SG-localized mRNAs undergo active translation. Related to Figure 4. (A) Excerpts from a time-lapse movie showing the buildup of SunTag signal on a SG-localized ATF4-SunTag mRNA (arrows). Cells were treated with 100 μM sodium arsenite for 1 h and imaged with 10 s intervals and Z-stack acquisition. Scale bar indicates 2 μm and time is shown in s. (B) Quantification of MS2 signal fluctuations on 11 mRNAs imaged with 10 s intervals and Z-stack acquisition. Black line indicates mean values. Same dataset as in Figure 4. (C) Excerpts from a time-lapse movie showing the loss of SunTag signal on a SG-localized ATF4-SunTag mRNA (arrows) after treatment with puromycin. After 1 h treatment with 100 μM sodium arsenite, 100 μg/mL puromycin was added and cells were immediately imaged. Scale bar indicates 2 μm and time is shown in s.

**Figure S4.**
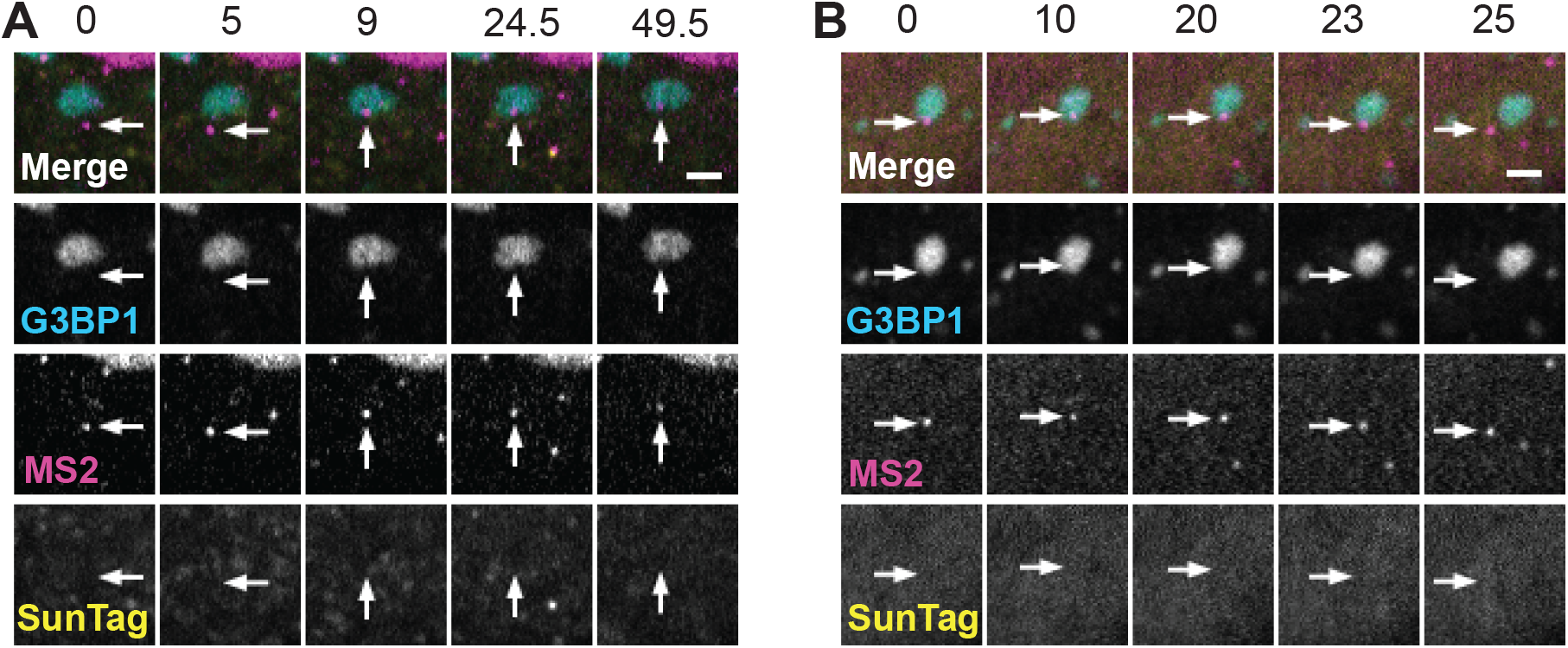
Non-translating mRNAs transition between cytosol and SGs. Related to Figure 5. (A) Excerpts from a time-lapse movie showing a non-translating ATF4-SunTag mRNA (arrows) arriving from cytosol to a SG. Cells were treated with 100 μM sodium arsenite for 1 h. Scale bar indicates 2 μm and time is shown in s. (B) Excerpts from a time-lapse movie showing a non-translating ATF4-SunTag mRNA (arrows) leaving a SG. Cells were treated with 100 μM sodium arsenite for 1 h. Scale bar indicates 2 μm and time is shown in s.

## SUPPLEMENTAL VIDEOS

**Video S1. Translating mRNAs localized to SGs.**

Example of SG-associated translation in cells treated with 100 μM sodium arsenite for 1 h. Arrows indicate ATF4-SunTag mRNAs labelled with SunTag signal and localized to SGs.

**Video S2. Long-lasting localization of translating mRNA to SG.**

Example of a long-lasting localization of SunTag-positive ATF4-SunTag mRNA to SG (arrow). Cells were treated with 100 μM sodium arsenite for 1 h, time-lapse movie was acquired with 10 s intervals and Z-stack acquisition.

**Video S3. SunTag signal buildup on SG-localized mRNA.**

Time-lapse movie showing the buildup of SunTag signal on a SG-localized ATF4-SunTag mRNA (arrow). Cells were treated with 100 μM sodium arsenite for 1 h.

**Video S4. Loss of SunTag signal after puromycin treatment.**

Time-lapse movie showing the loss of SunTag signal on a SG-localized ATF4-SunTag mRNA (arrow) after puromycin treatment. After 1 h treatment with 100 μM sodium arsenite, 100 μg/mL puromycin was added and cells were immediately imaged. Movie is paused at 25 s, showing the mRNA devoid of SunTag signal. After the pause, the mRNA remains localized to the SG.

**Video S5. Translating mRNA arriving to a SGs.**

Time-lapse movie showing a translating ATF4-SunTag mRNA (arrow) arriving from cytosol to a SG. Cells were treated with 100 μM sodium arsenite for 1 h.

**Video S6. Translating mRNAs leaving a SGs.**

Time-lapse movie showing a translating ATF4-SunTag mRNA (arrow) leaving a SG. Cells were treated with 100 μM sodium arsenite for 1 h.

**Video S7. SG-associated translation of 5’TOP mRNA.**

Example of SG-associated translation of the 5’TOP-SunTag mRNA in cells treated with 100 μM sodium arsenite for 50 min. SunTag-positive mRNA is indicated by an arrow.

**Video S8. ATF4-SunTag mRNA interacts transiently with a PB.**

Movie showing a translating ATF4-SunTag mRNA (arrow) transiently touching a PB in cells treated with 100 μM sodium arsenite for 1 h.

